# Immune phenotype-guided identification of disease-associated pathobionts in Crohn’s disease

**DOI:** 10.1101/2025.04.01.646605

**Authors:** Hiroko Nagao-Kitamoto, Kohei Sugihara, Sho Kitamoto, Shinya Ebihara, Yeji Kim, Kira L. Newman, Uxia Alonso, Takehiro Suzuki, Yoshiyuki Kioi, Arash Yunesi, Naohiro Inohara, Christopher J. Alteri, Phillip Minar, Lee A. Denson, Peter D. R. Higgins, John Y. Kao, Shrinivas Bishu, Yuying Xie, Yu Leo Lei, Nobuhiko Kamada

## Abstract

Aberrant immune activation within the gut mucosa and gut dysbiosis have been implicated in the pathogenesis of Crohn’s disease (CD). However, the specific immune responses triggered by dysbiotic microbiota, as well as the bacteria responsible for this activation, remain incompletely understood. Here, using the human microbiota-associated (HMA) mouse system, we demonstrated that colonization with dysbiotic gut microbiota from CD patients specifically induces the accumulation of mononuclear phagocytes, which may drive an interleukin-1 (IL-1)-driven inflammatory signature. Moreover, we identified pathobiont strains with a potent IL-1β-inducing capacity, termed ‘IL-1β-inducing pathobionts’ (IBIP). Isolated IBIP strains exhibit genetic and functional similarities to adherent-invasive *Escherichia coli* but harbor unique virulence-associated genes. Colonization with the IBIP *E. coli* strain exacerbated experimental colitis in an IL-1 signal-dependent manner. Notably, the colonization of IBIP *E. coli* can be detected by measuring the levels of specific immunoglobulin A (IgA) in their stool samples. Moreover, the level of IBIP-reactive IgA in stool may serve as a predictive biomarker for treatment response to anti-TNF therapies in treatment-naïve pediatric CD patients. Altogether, IBIP colonization could help identify CD patients with inflammatory dysbiosis who are likely to be refractory to anti-TNF therapies.

## Introduction

Inflammatory bowel disease (IBD), categorized into ulcerative colitis (UC) and Crohn’s disease (CD), is characterized by chronic and relapsing inflammation of the gastrointestinal tract. Although the precise etiology of IBD is not fully understood, various genetic, epigenetic, immunological, and environmental factors have been linked to its risk(1–3). Notably, growing evidence suggests that the gut microbiota plays a crucial role in the pathogenesis of IBD, both UC and CD(4–7). IBD patients exhibit an imbalanced composition of gut microbiota, so-called gut dysbiosis, characterized by an enrichment of potentially harmful commensal members at the expense of beneficial microbes, compared to healthy individuals (8, 9). This gut dysbiosis in IBD does not appear to be merely a secondary consequence of inflammation or medication effects. Instead, recent studies suggest a causal role for gut dysbiosis in the development and progression of intestinal inflammation. For example, the transplantation of microbiotas from IBD patients into germ-free mice (known as the human microbiota-associated (HMA) mouse model) triggers the development of spontaneous colitis in genetically susceptible hosts(7). Importantly, not all patient-derived microbiotas possess colitogenic capacity despite compositional imbalance(7). This suggests that the enrichment of specific pathobionts may confer the colitogenic trait to the microbiota in certain populations of IBD patients. In addition to the gut dysbiosis, dysregulated immune responses in the gut mucosal tissues are hallmarks of IBD. Recent bulk and single-cell transcriptomic analyses of gut tissue samples have highlighted distinctive cellular and immunological features associated with IBD, some of which are specific to particular disease types (i.e., CD or UC) or correlate with disease characteristics, such as location, activity, and response to treatment(10–14). However, the precise immune responses triggered by the dysbiotic microbiota, along with the specific bacteria responsible for initiating this activation, are still not fully understood.

Here, using the HMA mouse model, we demonstrated that colonization with dysbiotic gut microbiota from CD patients specifically induces the accumulation of *Il1b*^hi^ mononuclear phagocytes. This accumulation may trigger an IL-1-driven inflammatory signature observed in human IBD patients, which is associated with histological inflammation, ulceration, and treatment nonresponse(12). Moreover, we identified specific pathobionts responsible for eliciting robust IL-1β production by mononuclear phagocytes, which we termed ‘IL-1β-inducing pathobionts’ (IBIP). The IBIP strains we isolated share genetic and functional similarities with known adherent-invasive *E. coli* (AIEC) strains, yet they also harbor virulence genes associated with extraintestinal pathogenic *E. coli* (ExPEC). Colonization with the IBIP *E. coli* strain exacerbated experimental murine colitis in an IL-1 signal-dependent manner. Also, the abundance of IBIPs in stool samples, measured by IBIP-reactive IgA levels, may help assess the colitogenic potential of gut microbiota in CD. Notably, high levels of IBIP colonization, as indicated by fecal IgA levels, are likely associated with treatment failure of anti-TNF therapies in treatment-naïve pediatric CD patients. Collectively, IBIP colonization may serve as a biomarker for identifying CD patients at risk of severe histological inflammation or nonresponse to treatment. Targeted therapies aimed at decolonizing these bacteria could provide significant benefits for these patients.

## Results

### IL-1β is a central mediator of CD microbiota-associated colitis

To identify disease-driving pathobionts in CD patients, we first examined which immune pathways are selectively activated by the microbiota from CD patients but not healthy control (HC) subjects. To this end, we employed the HMA mouse model, in which GF *Il10*^−/-^ mice were colonized by the microbiotas either from CD or HC (7). Stool samples were collected from CD patients and HC subjects (5 different individuals each) and transplanted into GF *Il10*^−/-^ mice (n=2 per individual) and maintained for 3 weeks (**Figure 1A** and **Table S1**). Consistent with previous studies (7), some key features of CD donor microbiotas, such as the decreased Shannon α-diversity index, were successfully recapitulated in CD-HMA mice (**Figure S1A-G**). To examine ethe impact of CD dysbiotic microbiota on immune activation in the colon, the colonic lamina propria (LP) cells were isolated from HC-and CD-HMA mice and subjected to immune profiling by single-cell RNA sequencing (scRNA-Seq) (**Figure 1A**). We identified 21 clusters defined by gene expression patterns (**Figure 1B,C** and **Figure S2**). Among these clusters, only 2 clusters (clusters 3 and 19) were significantly expanded in CD-HMA mice compared to HC-HMA mice (**Figure 1D**). Clusters 3 and 19 express high levels of *Cd14*, *Itgam*, *Itgax*, *Tnf*, *Il23a*, and *Osm*, indicating that they represent inflammatory mononuclear phagocytes (MNPs) (**Figure 1C**). Moreover, cluster 3 expresses *Lyz2*, while cluster 19 expresses *Mrc1* and *Cd274*, suggesting that cluster 3 is macrophage-like, whereas cluster 19 resembles DC-like MNPs (**Figure 1C**). Notably, these MNP clusters express high levels of IL-1 signaling-related genes, including *Il1b*, *Il1a*, *Nlrp3*, and *Casp1*, suggesting that they are major producers of IL-1α and IL-1β in the colonic mucosa (**Figure 1C**). Thus, the observed expansion of these MNPs in CD-HMA mice may play a key role in initiating IL-1-mediated inflammatory responses in the colon. Consistent with this notion, gut inflammation induced by the colonization of CD microbiotas in *Il10*^−/-^ HMA mice was significantly attenuated by the treatment with the IL-1R antagonist Anakinra (**Figure 1E-G**). These data suggest that IL-1-expressing MNPs are a crucial mediator of CD microbiota-driven colitis in HMA mice.

**Figure 1.**
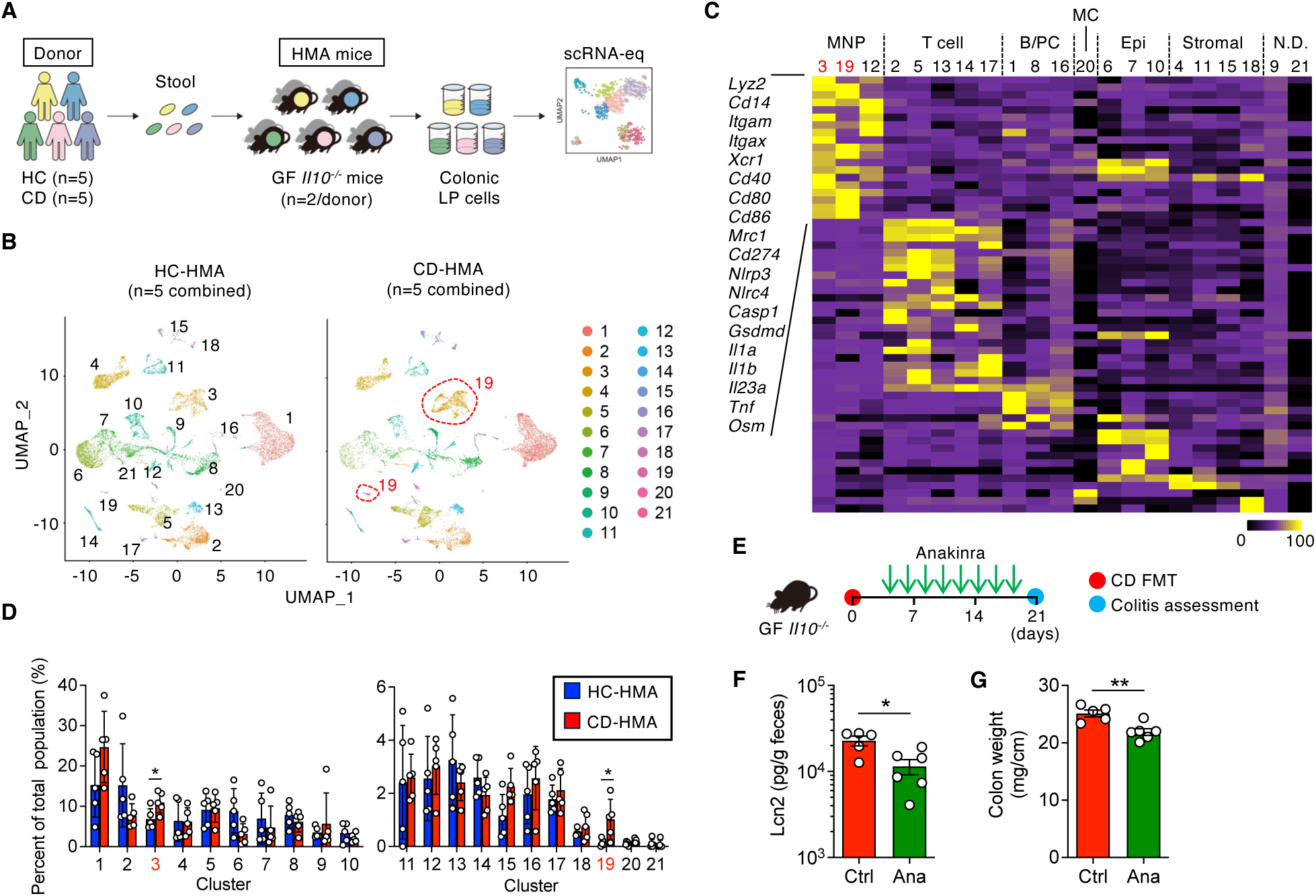
Activation of IL-1β-producing mononuclear phagocytes as key mediators of inflammation in the CD microbiota-driven colitis model. (A) Stool samples were collected from five healthy control (HC) subjects and five patients with Crohn’s disease (CD). Fecal microbes were inoculated into germ-free (GF) *Il10^−/-^* mice to generate human microbiota-associated (HMA) mice (two mice per donor). After three weeks of bacterial reconstitution, the cecum and colon tissues were harvested, and lamina propria (LP) cells were isolated for single-cell RNA sequencing analysis. (B) Uniform manifold approximation and projection (UMAP) plot displaying HC-HMA and CD-HMA mice, with cells colored by cluster type. (C) Relative expression levels of the indicated genes across 21 cell clusters. (D) Proportional abundance of each cell cluster. (E) CD-HMA *Il10^−/-^* mice colonized with microbiota from CD005 were treated with either PBS (control) or Anakinra eight times following microbiota colonization. Mice were sacrificed on day 21. (F) Fecal lipocalin-2 (Lcn2) levels on day 21. (G) Colonic weight. (F, G) Data are presented as mean ± SD, with dots representing individual mice (*n* = 5–6). *; *P* < 0.05, **; P < 0.01 (Mann-Whitney *U* test).

### Isolation of IL-1β-inducing pathobionts from CD microbiota

Next, guided by the identified immune profiles, we aimed to identify specific pathobionts within CD microbiotas that contribute to colitis induction. To this end, we focused on bacterial strains capable of inducing IL-1β secretion from MNPs. We harvested gut bacteria from CD-HMA mice and cultured them in various bacteria growth mediums under anaerobic conditions. Several bacterial colonies were isolated and examined for their ability to induce IL-1β production in bone marrow-derived macrophages (BMDMs). Among these, only a few bacterial strains from CD-HMA mice exhibited a potent IL-1β-inducing capacity (**Figure 2A and 2B**). Interestingly, these strains did not stimulate the secretion of the anti-inflammatory cytokine IL-10, suggesting that they function as potential pathobionts that trigger inflammatory responses (**Figure 2A and 2B**). We identified three bacterial strains - HK1, HK2, and HK7 - that selectively induced IL-1β but not IL-10, classifying them as IL-1β-inducing pathobionts (IBIPs). Notably, these IBIP strains were absent in HC-HMA mice, indicating their likely enrichment in CD patients (**Figure 2A and 2B**). In contrast, bacteria capable of inducing tumor necrosis factor (TNF) were found in both HC-HMA and CD-HMA mice, with some of these strains also stimulating high levels of IL-10 (**Figure 2A and 2B**). The genomic analysis identified that isolated IBIPs HK1, HK2, and HK7 are *Escherichia coli* (*E. coli*) and exhibit genetic similarities with IBD-associated adherent-invasive *E. coli* (AIEC) strains (**Figure 2C**) (15, 16). However, IBIP strains lack certain AIEC-associated virulence genes, such as *lpfA* and *pduC* (**Figure 2D** and **Figure S3**). Notably, isolated IBIP *E. coli* strains possess virulence genes associated with extraintestinal pathogenic *E. coli* (ExPEC), such as *pap*, *cnf1*, and *senB*, which are not present in reference AIEC strains such as LF82, CUMT8, and NC101 (**Figure 2D** and **Figure S3**). Further, the IBIP *E. coli* strains belong to the B2 phylogroup **(Figure 2C**); the majority of ExPEC isolates are found within B2. Additionally, these strains possess the K1 capsule serotype that are generally restricted to uropathogenic and neonatal meningitis ExPEC isolates. Next, we analyzed functional characteristics of isolated IBIP strains. Since IBIP *E. coli* HK1 and HK2 are almost genetically identical, we used HK1 and HK7 strains in subsequent analyses. Consistent with the genetic features, IBIP *E. coli* HK1 and HK7 strains and strong adherence and invasion phenotype when cultured with colonic epithelial cells (**Figure 2E**). Next, we validated the extent to which IBIP *E. coli* HK1 and HK7 strains induce the secretion of IL-1β by the colonic cells. Consistent with BMDMs, colonic lamina propria (LP) cells isolated from healthy mice secreted IL-1β upon stimulation with IBIP *E. coli* HK1 and HK7, but not with non-pathogenic commensal or probiotic *E. coli* strains (HS, MG1655 [MG], and Nissle1917 [EcN]) and AIEC strains (LF82 and CUMT8) (**Figure 2F**). In contrast, IL-10 induction by IBIP *E. coli* strains was significantly weaker than that triggered by the other *E. coli* strains. In LP cells from mice with DSS-induced colitis, IBIP *E. coli* strains elicited a robust IL-1β response, whereas non-pathogenic and AIEC strains induced only minimal IL-1β, even in the inflamed environment. Similar to healthy LP cells, IL-10 induction by IBIP strains in inflamed LP cells remained significantly lower than that of other *E. coli* strains. As a result, the IL-1β/IL-10 ratio induced by IBIP *E. coli* strains in inflamed LP cells was markedly higher than that observed with commensal or AIEC strains (**Figure 2G**). In contrast to IL-1β and IL-10, TNF induction was comparable across all *E. coli* strains tested (**Figure 2F**). IL-1β, but not TNF, induced by IBIP stimulation was diminished in *Nlrp3^−/-^*, *Pycard*^−/-^, and *Casp1^−/-^* BMDMs, indicating that IBIPs activate the Nlrp3 inflammasome (**Figure S4**). Nlrp6 may also be partially involved in IL-1β by IBIPs, especially strain HK7 (**Figure S4**). The lack of *Myd88* abolished the secretion of both IL-1β and TNF, suggesting that TLR signaling is activated by IBIP stimulation. Thus, isolated pathobiont strains responsible for the induction of immune activation signatures in CD-HMA mice share genetic and functional similarities with IBD-associated but may harbor unique virulence traits.

**Figure 2.**
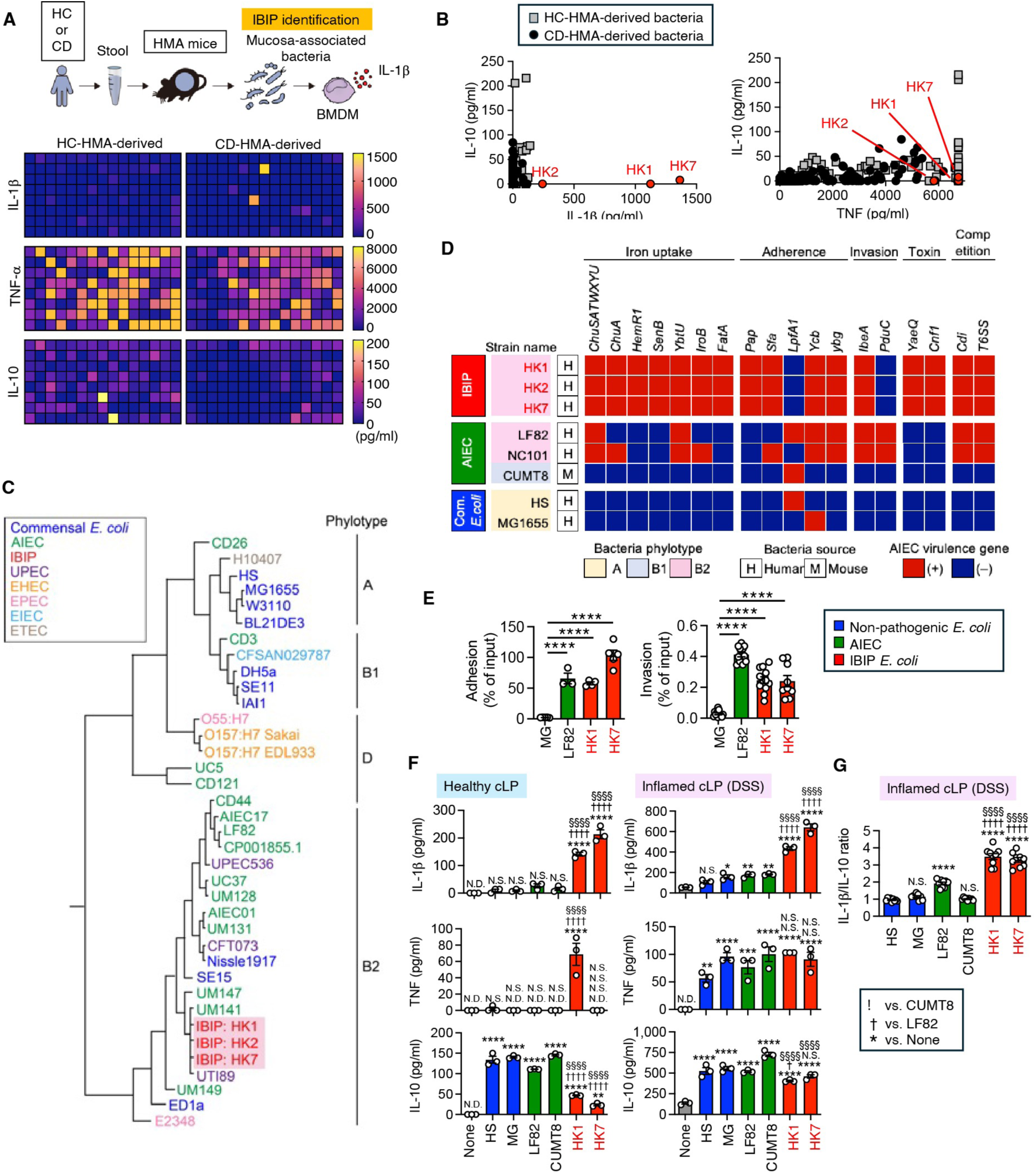
Identification of IL-1β-inducing pathobionts residing in CD microbiotas. (A) Mucosa-associated bacteria were isolated from colonic tissues of HC-HMA and CD-HMA mice. The isolated bacteria were used to stimulate BMDMs, and cytokine production was measured. Cytokine levels are presented as heatmaps. (B) Correlation analysis between the pro-inflammatory cytokines IL-1β and TNF and the anti-inflammatory cytokine IL-10 induced by each bacterial isolate. Red dots indicated the IBIP strains isolated from CD. (C) Phylogenetic tree of the identified IL-1β-inducing pathobiont (IBIP) strains (HK1, HK2, and HK7). (D) Presence (red) or absence (blue) of AIEC and ExPEC-associated virulence genes in IBIP strains, commonly studied AIEC strains (LF82, NC101, CUMT8), and commensal *E. coli* strains (HS, MG1655). (E) [Left] T84 cells were infected with commensal *E. coli* (MG1655), AIEC (LF82), and IBIP strains (HK1 and HK7) at a MOI of 10. After 3 hours, cells were lysed and plated to determine CFUs, representing the total number of adherent bacteria. [Right] To assess intracellular bacterial load, gentamicin was added for 1 hour to eliminate extracellular bacteria, and lysed cells were plated. Data are presented as mean ± SD (*n* = 3–12). **** *P* < 0.0001 by one-way ANOVA followed by Bonferroni post hoc test. (F and G) (F) Colonic lamina propria (cLP) cells from healthy or inflamed (DSS colitis) mice were stimulated with commensal *E. coli* (HS and MG1655), AIEC (LF82 and CUMT8), and IBIP strains (HK1 and HK7). Levels of IL-1β, TNF, and IL-10 were measured in culture supernatants. (G) The ratio of IL-1b and IL-10. Data are presented as mean ± SD, with dots representing individual mice (*n* = 3-9). *^, †^; P < 0.05, **; P < 0.01, ***; P < 0.001, ****^, ††††, §§§§^; P < 0.0001, by 1-Way ANOVA followed by Bonferroni post hoc test. N.D.: Not detected. N.S.: Not significant.

### IBIP colonization aggravates an animal model of IBD

We next examined the sufficiency of IBIP colonization in the exacerbation of colitis. To this end, SPF mice were stably colonized by the IBIP HK7 strain or a control non-IBIP commensal *E. coli* strain MG1655 by multiple gavage. Mice were then kept for 21 days to allow enough time to modulate immune activation by IBIP colonization. After 21 days of *E. coli* colonization, mice were then given DSS to trigger colitis (**Figure 3A**). To assess the involvement of IL-1 signal activation by IBIP HK7 in the pathogenesis of colitis, one group of mice was treated with anakinra (**Figure 3A**). The levels of colonization of *E. coli* were comparable between groups (i.e., MG1655, IBIP HK7, and HK7 + anakinra) during pre-colonization (**Figure 3B**). The IBIP HK7 without anakinra group showed higher *E. coli* colonization levels on days 24 and 28 (i.e., 3 and 7 days after DSS administration), likely due to a greater degree of inflammation compared to the other groups. (**Figure 3B**). All mice were euthanized on day 28 (7 days post-DSS), and the severity of colitis was evaluated. As a result, mice colonized with IBIP exhibited more severe colitis, as seen in the shortening and thickening of the colon, histological inflammation, and pro-inflammatory gene expression in the colonic tissues, compared with the control groups (i.e., mock infection and MG1655 colonization) (**Figure 3C-F**). Notably, the severity of IBIP-induced colitis phenotypes was significantly attenuated by the anakinra treatment, despite IBIP colonization was not affected by anakinra. These results suggest that IBIP colonization exacerbates DSS-induced colitis through the activation of IL-1 signaling.

**Figure 3.**
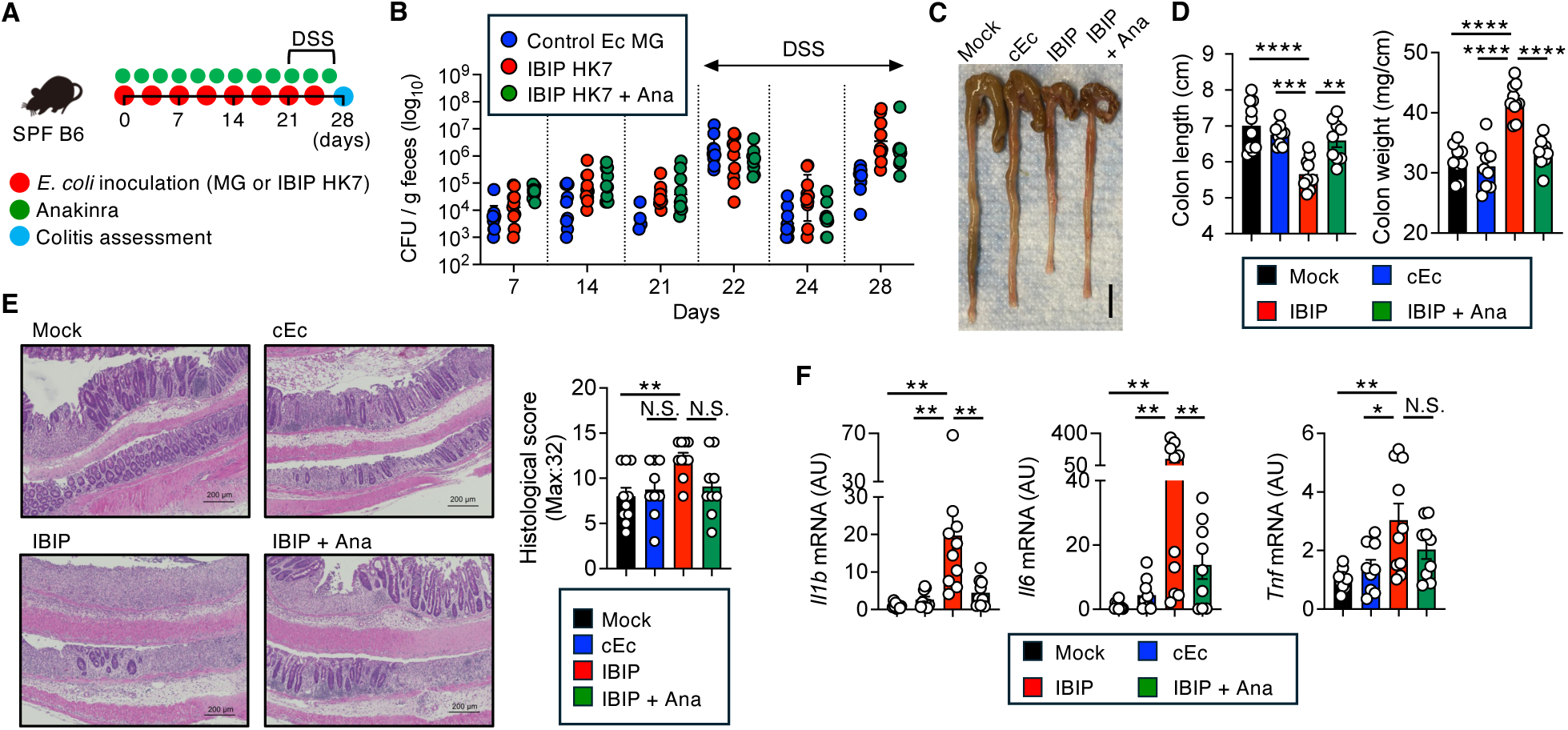
IBIP colonization exacerbates colitis in an animal model of IBD. (A) SPF-B6 mice were pre-treated with streptomycin and ampicillin and subsequently colonized with either control *E. coli* (MG1655) or the IBIP *E. coli* (HK7) (inoculated twice a week for 4 weeks). The mock colonization group received PBS gavage. An IBIP-colonized group was treated with Anakinra 3 times a week for 4 weeks (IBIP + Ana). Twenty-one days after the first bacterial inoculation, all mice were administered 1.5% DSS for five days, followed by two days of water. (B) *E. coli* CFUs in fecal samples. (C-G) On day 28, mice were euthanized, and colitis severity was assessed. (C) Representative macroscopic images of the cecum and colon. (D) Colon length and weight. (E) [Left] Representative histological images. The scale bar is 200 μm. [Right] Histological score of colonic inflammation. (F) mRNA expression of pro-inflammatory cytokines in the colonic tissues. Data are presented as mean ± SD. Dots represent individual mice. Bars represent the median in (B). Data were pooled from 2 independent experiments (total n =10). N.S.: Not significant. *; P < 0.05, **; *P* < 0.01, ***; P < 0.001, ****; P < 0.0001 by 1-Way ANOVA followed by Bonferroni post hoc test.

### IBIP colonization may serve as a biomarker for predicting the colitogenic capacity of the gut microbiota and patients’ treatment response

Next, we aimed to determine the prevalence of IBIP colonization in CD patients. Although we identified IBIP *E. coli* strains carrying ExPEC virulence genes, the specificity of these genes in detecting IBIP colonization remains uncertain. To address this, we assessed luminal immunoglobulin A (IgA) levels against IBIPs, as gut bacteria with colitogenic potential are typically highly coated with IgA (17), providing an alternative biochemical measure. We analyzed stool samples from HC subjects (n=7) and CD patients (n=18) (**Table S1**), measuring the levels of luminal IgA reactive to IBIP HK7. The mean + 3 S.D. value from HC samples was used as a cut-off, and individuals with IgA concentrations above this threshold were classified as having high IBIP HK7-reactive IgA levels (**Figure 4A)**. As a result, 10 out of 19 CD patients (55%) tested positive for IBIP HK7-reactive IgA, indicating that approximately half of the patients had or have been colonized with IBIPs. IBIP HK7-reactive IgA levels did not correlate with disease activity, suggesting that increased IBIP colonization is not solely a consequence of inflammation (**Figure 4B**). Next, we sought to determine whether IBIP HK7 colonization serves as a biomarker for predicting the colitogenicity of dysbiotic microbiota in CD. To this end, GF *Il10*^−/-^ mice were colonized with microbiota from HC (n=7) and CD (n=11) donors. The relationship between fecal IBIP HK7-reactive IgA levels in donor stools and colitogenicity in *Il10*^−/-^ HMA mice, as measured by fecal Lcn2 level, was examined (**Figure 4C**). Consistent with our previous study, none of the HC donor stool samples induced colitis in *Il10*^−/-^ HMA mice (**Figure 4C**) (7). In contrast, 4 out of 11 (36%) tested CD donor stools induced colitis in *Il10*^−/-^ HMA mice, all of which were positive for IBIP HK7-reactive IgA (**Figure 4C**). However, 2 CD microbiotas with high IBIP HK7-reactive c IgA levels did not exhibit colitogenic capacity (**Figure 4C**). Similar to HC microbiotas, none of the CD microbiotas negative for IBIP HK7-reactive IgA displayed colitogenicity in *Il10*^−/-^ HMA mice (**Figure 4C**). Notably, we observed the same phenotype when IBIP HK1 strain-reactive IgA was used as a biomarker, suggesting that these IgAs recognize common antigens present in IBIP strains, which may serve as determinants of their functional traits (**Figure S5**). Thus, IBIP colonization, as measured by IBIP-reactive IgA levels in stool samples, may be a characteristic indicator of the colitogenic potential of dysbiotic microbiota in CD patients, although this is not the only factor rendering microbiota to be colitogenic.

**Figure 4.**
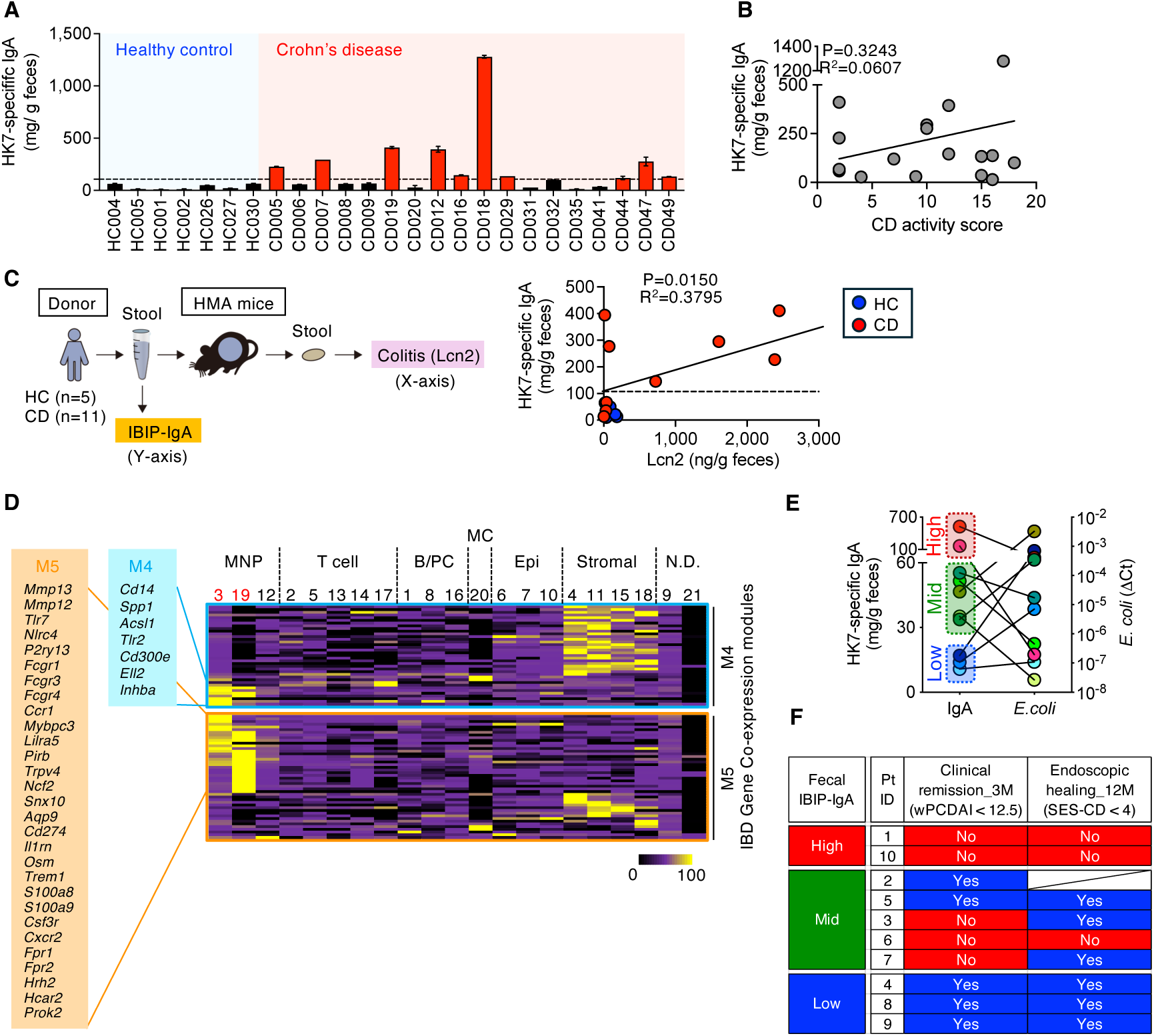
IBIP-reactive IgA as a potential biomarker for colitogenic microbiota and treatment response in CD patients. (A) Levels of IBIP HK7-specific IgA in fecal samples from 7 HC and 18 CD patients. Data are presented as mean ± SD (*n* = 2 technical replicates). The dotted line represents the cut-off value (mean + 3 SD of HC samples). (B) Correlation between IBIP HK7-reactive IgA levels and CD activity scores (Harvey-Bradshaw Index, HBI). Statistical significance was determined using Pearson correlation analysis (two-sided). (C) Correlation between IBIP HK7-reactive IgA levels and the colitogenic capacity of microbiotas. The colitogenic potential of microbiotas was assessed by colonizing GF *Il10^−/-^* mice (HMA mice). After 3 weeks of bacterial reconstitution, fecal Lcn2 levels were measured to determine the degree of colitis. Statistical significance was determined using Pearson correlation analysis (two-sided). (D) Heatmap showing the expression of genes in modules M4 and M5, which are specifically associated with treatment nonresponse, across different clusters in the scRNA-seq analysis from Figure 1. (E) Correlation between IBIP HK7-reactive IgA levels and *E. coli* abundance (measured by qPCR) in treatment-naïve pediatric CD patients at baseline. (F) Pediatric CD patients from (E) received anti-inflammatory therapies. The relationship between baseline IBIP HK7-reactive IgA levels and treatment response is shown. Clinical remission was defined as weighted Pediatric Crohn’s Disease Activity Index (wPCDAI) < 12.5 at 3 months post-treatment. Endoscopic healing was determined by a Simple Endoscopic Score for CD (SES-CD < 4) at 12 months post-treatment.

A previous study reported that the accumulation of IL-1β-expressing MNPs and following activation of stromal cells and recruitment of neutrophils are a known hallmark of CD patients who are non-responders to various therapies, including anti-TNF therapies and corticosteroids(10, 12, 18). In this regard, the previous study identified 38 gene co-expression modules (M1 – M38) in IBD patients, with modules M4 and M5 specifically associated with treatment nonresponse(12). Notably, we found that several genes within these modules—such as *Cd14*, *Osm*, *S100a8/9*, *Mmp12/13*, and *Il1rn*—are expressed in MNP clusters 3 and 19, which were expanded in CD microbiota colonized HMA mice (**Figure 4D**). Thus, it is plausible that IBIP colonization drives an immune cell network signature characterized by M4/M5 modules through IL-1β induction, thereby contributing to treatment refractoriness. To test this hypothesis, we measured baseline IBIP-reactive IgA levels in stool samples from treatment-naïve pediatric CD patients and evaluated the relationship between IBIP colonization and response to anti-TNF therapies (**Table S2**). Among the 10 pediatric CD patients, 2 exhibited high IBIP-specific IgA levels in stool samples (**Figure 4E**). Additionally, 5 patients had medium levels, while the remaining 3 displayed low IBIP-reactive IgA levels (**Figure 4E**). Notably, IBIP-specific IgA levels did not correlate with *E. coli* abundance in stool, as quantified by qPCR. This suggests that increased IBIP-specific IgA is not merely a reflection of higher *E. coli* colonization but rather indicates selective colonization by IBIPs (**Figure 4E**). Patients’ response to anti-TNF therapies (infliximab or adalimumab) was evaluated based on clinical remission (wPCDAI < 12.5) at 3 months post-treatment and endoscopic healing (SES-CD < 4) at 12 months post-treatment. Intriguingly, patients with high baseline IBIP-reactive IgA levels were refractory to anti-TNF therapies, whereas those with low IBIP-reactive IgA levels responded to treatment (**Figure 4F**). Patients with mid-range IBIP-reactive IgA levels included both responders and non-responders (**Figure 4F**). These findings suggest that IBIP colonization could serve as a predictive marker for therapeutic response in pediatric CD patients.

## Discussion

By integrating humanized gnotobiotic animal technology with scRNA-Seq, we identified host gene transcriptional signatures induced by the colonization of dysbiotic microbiota in CD patients. The gene expression pattern triggered by CD-associated microbiota is characterized by the accumulation of *Il1b*⁺ MNPs and a subsequent IL-1-driven inflammatory signature, which has been previously reported to be associated with histological inflammation, ulceration, and resistance to treatments such as anti-TNF therapies (12). Pathobionts capable of inducing robust IL-1β secretion by MNPs, termed IL-1β-inducing pathobionts (IBIP), have been identified as key bacteria responsible for eliciting such immune responses. Notably, colonization by IBIP is sufficient to exacerbate colitis in the DSS-induced IBD model.

Several pathogenic microorganisms, such as *Salmonella enterica* serovar Typhimurium, are known to activate the inflammasome and trigger the release of bioactive IL-1β. In contrast, the normal commensal microbiota fails to induce inflammasome-IL-1 signaling(19–21). In this context, some studies have shown that certain commensal pathobionts - typically minor constituents of the healthy gut microbiota but accumulating under specific disease conditions - can activate inflammasomes and cleave pro-IL-1β, leading to the secretion of mature IL-1β by intestinal MNPs in mice(21, 22). For example, *Proteus mirabilis*, which expands during colitis in SPF mice, activates the NLRP3 inflammasome and induces IL-1β production through hemolysin(21). Similarly, oral commensal *Klebsiella* and *Enterobacter* species, such as *K. aerogenes*, stimulate IL-1β production by intestinal MNPs when they ectopically colonize the gut mucosa(22). Notably, these abnormally expanded (IBIPs play pivotal roles in triggering and/or exacerbating colitis through inflammasome-IL-1 signaling in various IBD mouse models(21, 22). However, in human IBD, the precise identity of disease-driving pathobionts capable of activating inflammasome-IL-1 signaling remains largely unexplored. In this study, we identified *E. coli* strains harboring genetic and functional characteristics of AIEC and ExPEC as IBIPs residing in the human gut.

Notably, IBIP *E. coli* strains are not merely “immunogenic,” as they do not induce the anti-inflammatory cytokine IL-10. Many latent pathobionts can stimulate both pro-inflammatory and anti-inflammatory cytokines in macrophages, thereby contributing to gut homeostasis in immunocompetent hosts. However, their colonization leads to excessive immune activation in immunocompromised hosts, such as Rag- or IL-10-deficient mice. For example, *Helicobacter hepaticus* induces an anti-inflammatory gene signature, including IL-10, in macrophages, making it non-pathogenic in wild-type hosts(23). In contrast, *H. hepaticus* colonization leads to severe colitis in IL-10-deficient hosts (24). Similarly, colonization with a CD patient-derived AIEC strain exacerbates DSS-induced colitis only in the absence of IL-10(25). In this context, IBIP *E. coli* strains aggravated DSS-induced colitis even in IL-10-sufficient wild-type mice. This may be due to their ability to selectively induce pro-inflammatory cytokines, such as IL-1β, without simultaneously inducing anti-inflammatory cytokines. Thus, IBIP *E. coli* strains may contribute to inflammation in IBD patients without genetic or immunological defects in IL-10 signaling pathways.

The IL-1β-inducing capacity of IBIP *E. coli* strains is significantly stronger than that of commonly studied AIEC strains, such as LF82. This suggests that virulence factors unique to IBIP *E. coli* strains may be responsible for activating the inflammasome-IL-1 axis. In this context, certain virulence genes found in IBIP *E. coli* strains but absent in reference AIEC strains (LF82, NC101, and CUMT8) are suspected to trigger inflammasome activation. For instance, cytotoxin necrotizing factor-1 (CNF1), a virulence factor in uropathogenic *E. coli*, has been shown to activate the NLRP3 inflammasome (26). Also, HlyC, an internal protein acyltransferase, activates the *α*-hemolysin toxin(27, 28), and active *α*-hemolysin induces NLRP3-mediated IL-1 secretion(21). These virulence genes may, therefore, contribute to the robust IL-1β induction observed in IBIP *E. coli* strains. Further studies are needed to identify the specific virulence factors responsible for inflammasome activation in IBIP *E. coli* strains.

While the pathogenic role of IL-1β in IBD is widely recognized, some conflicting evidence remains. Numerous studies have reported that IL-1β expression is significantly elevated in the intestinal mucosa of patients with CD compared to healthy controls(29). Moreover, genetic variations in *NLRP3*, an inflammasome protein associated with the secretion of bioactive IL-1β, have been linked to an increased risk of CD(30, 31). Together, these findings suggest that excessive activation of the NLRP3 inflammasome-IL-1β pathway plays a critical role in IBD pathogenesis. Supporting this notion, IL-1 signaling blockade has been shown to ameliorate intestinal inflammation in several preclinical models of IBD(21, 22, 32, 33). In human IBD, while large-cohort clinical studies are lacking, small-cohort studies - particularly in IL10R-deficient very early-onset (VEO)-IBD patients - have demonstrated the efficacy and safety of therapies targeting IL-1 signaling(33, 34). Consistent with this, several agents that block NLRP3-IL-1 signaling are currently under clinical evaluation(35). However, some studies have also shown that NLRP3 and IL-1β contribute positively to epithelial barrier integrity (36, 37). Therefore, therapies targeting the NLRP3-IL-1 signaling pathway may not be beneficial for all IBD patients. In this regard, our study found that IBIP colonization levels vary among patients. It is possible that patients with high IBIP colonization may respond better to therapies targeting IL-1 signaling. Supporting this idea, IBIP colonization levels may serve as a biomarker for predicting treatment response in IBD patients. Recent studies have demonstrated that enrichment of IL-1β-expressing MNPs is associated with resistance to corticosteroid and anti-TNF treatment in IBD patients (10, 12, 18). Indeed, our pilot study in a small cohort of treatment-naïve pediatric IBD patients suggests that IBIP colonization, determined by fecal IgA levels, may predict treatment response to anti-TNF therapies. Further studies with larger patient cohorts are needed to verify the correlation between IBIP colonization and anti-TNF resistance. If confirmed, high IBIP colonization could serve as an effective biomarker for predicting response to anti-TNF therapies. Moreover, these patients may benefit more from IL-1-targeted therapies than from anti-TNFs.

In conclusion, we have successfully identified pathobionts that drive an IBD-associated immune gene signature using combined gnotobiotic and single-cell immune profiling technologies. These pathobionts represent potential therapeutic targets for mitigating the inflammatory potential of the gut microbiota in IBD patients. Furthermore, their colonization could serve as a biomarker for predicting treatment responses in IBD. By linking specific types of dysbiosis to immune alterations in IBD and identifying keystone pathobionts that drive these immune phenotypes, we may establish catalogs of pathobiont-immune activation combinations. Distinct pathobionts may be associated with resistance to different IBD therapies, and their colonization status could help guide the selection of the most effective biologic treatments for individual patients.

## Acknowledgments

The authors wish to thank the University of Michigan Microbiome Core, the Germ-Free Mouse Facility, the Advanced Genomics Core for technical assistance, Ingrid L. Bergin from the In-Vivo Animal Core at the University of Michigan Unit for Laboratory Animal Medicine for histological assessment, Merritt G. Gillilland III for the microbiome analysis, and Nana Iwami and Mikiko Kaiya for technical assistance. This work was supported by the National Institute of Health DK108901 (N.K.), DE026728 (Y.L.L.), JSPS KAKENHI JP24K22122 (H.N.-K), JP23H00404 (N.K.), JP23K17428 (S.K.), JP23K27586 (S.K.), JST FOREST Program JPMJFR220O (S.K.), Crohn’s and Colitis Foundation Research Fellowship Award and Career Development Award (H.N.-K. and K.S.), the Leona M. and Harry B. Helmsley Charitable Trust (L.A.D. and P.M).

## Authorship Contributions

H.N.-K., K.S., and N.K. conceived and designed experiments. H.N.-K. and K.S. conducted most of the experiments with help from S.K., S.E., Y.Kim., K.L.N., U.A., T.S, and Y.Kioi.. A.Y., Y.X., and Y.L.L analyzed scRNA sequencing data. N.I. performed microbiome and microbial genomic analyses. C.J.A. helped with critical advice on bacterial genomics. S.B., P.D.R.H., J.Y.K., P.M., and L.A.D. provided human stool samples and helped with critical advice and discussion. H.N.-K., K.S., and N.K. analyzed the data. H.N.-K. and N.K. wrote the manuscript with contributions from all authors.

## Data Availability

The microbial genomic data, microbiome data, and scRNA-Seq data presented in this study are available at the NCBI Sequence Read Archive under BioProject PRJNA1163383.

## Methods

### Sex as a biological variable

Our study included both male and female mice and patients, reporting similar findings across sexes.

### Mice

Specific pathogen-free (SPF) wild-type C57BL/6 mice were sourced from Jackson Laboratory and Japan SLC. The genetically altered mouse models, which included *Nlrp3*^−/-^, *Nlrc4*^−/-^, *Nlrp6*^−/-^, *Casp1*^−/-^, *Casp11*^−/-^, *Aim2*^−/-^, *Myd88*^−/-^, and *Pycard*^−/-^ strains, were generously supplied by Dr. Gabriel Núñez at the University of Michigan. All mice were bred and maintained within the Animal Facility at the University of Michigan or the University of Osaka, as previously described(22, 38). The experimental protocols were thoroughly reviewed and sanctioned by the Institutional Animal Care and Use Committee (IACUC) at both universities. All animal studies were performed in compliance with the ARRIVE guidelines and the ethical standards established by the University of Michigan and the University of Osaka.

### Human microbiota-associated gnotobiotic mice

Germ-free (GF) *II10^−/-^* mice were maintained in the Germ-Free Facility at the University of Michigan. GF mice were housed in flexible film isolators, and their GF status was monitored weekly through aerobic and anaerobic fecal cultures, along with microscopic examination of stained fecal contents to detect any unculturable contaminants. To establish human microbiota-associated (HMA) gnotobiotic mice, GF mice were colonized with stool samples collected from individuals with Crohn’s disease (CD) and healthy controls (HC) under protocols HUM00041845 and HUM00127476, approved by the University of Michigan Institutional Review Board. Written informed consent was obtained from all participants prior to the collection of samples. Both CD patients and HC individuals had not received any antibiotics for a minimum of three months before sample collection. Furthermore, the subjects had no reported history of intestinal bacterial infections, such as *Clostridioides difficile*, or other viral infections, including hepatitis B, hepatitis C, or human immunodeficiency virus. Patients with CD were diagnosed histologically and endoscopically prior to their inclusion in the study. The collected fecal samples were stored at -80°C until utilized. Before inoculation, the stool samples were diluted 1:10 in pre-reduced phosphate-buffered saline under anaerobic conditions. The diluted samples were then filtered through a 100 μm cell strainer to administer an oral inoculation of 100 μl per mouse to the GF *II10*^−/-^ mice. The HMA mice were housed in positive-pressure, individually ventilated cages (IVCs) (ISOcage P) to prevent cross-contamination and preserve their gnotobiotic condition(39, 40). All mice were provided with a sterilized rodent breeder diet (LabDiet 5013). Both male and female mice aged 8 to 16 weeks were utilized for the experiments, as previously describe(7, 38). The HMA *Il10^−/-^*mice were euthanized three weeks post-colonization with human microbiota. Fecal samples were collected for microbiome analysis and lipocalin-2 (Lcn2) measurements, while colonic tissues were used for cell isolation. For the Anakinra treatment, the HMA *Il10^−/-^* mice received intraperitoneal injections of Anakinra (Swedish Orphan Biovitrum AB) or saline every other day (50 mg/kg per mouse per dose) starting from day 4 after microbiota colonization until day 21.

### Single-cell RNA sequencing

We selected high-quality cells exhibiting unique feature counts ranging from 200 to 7,500, with mitochondrial read percentages below 20%. Cells with fewer than 500 unique RNA counts were excluded from the analysis. After the filtering process, a total of 28,599 high-quality cells were retained for further integration. The transform normalization process was conducted using Seurat v.3 to account for variations due to cell cycle effects, mitochondrial percentages, and library size. To enhance the segregation of immune lineages, we identified the top 3,000 highly variable genes alongside a selection of immune signature genes for integration based on Mutual Nearest Neighbors. The immune anchor genes included *Cxcr5, Cd69, Aim2, Irf5, Irf1, Irf3, Lgals9, Ly6e, Nos2, Il6, Tnf, Isg15, Gsdmd, Cd8a, Cd40, Cd80, Cd86, Il10, Tgfb1, Tmem173, Batf2, Cd274, Cxcl9, Mrc1, Siglec15, Trdc, Cd2, Trac, Cd4, Cd8b1, Foxp3, Trbc1, Trbc2, Gzmb, Eomes, Icos, Cd3d, Cd3e, Ifng, Ncr1, Cd19, Cd79a, Cd79b, Itgam, Itgax, Batf3, Xcr1, Gata3, Ctla4, Rorc, Il17a, Bcl6, Havcr2, Tnfrsf4, Tigit, Cxcl10, Mx1, Ifnb1, Il3ra, Nrp1, Fcer1a, Tbx21, Lag3, Ifnl3, Pdcd1, Cd14, Ifna4, Ly6g, Ly6c1*, and *Lyz2*. Utilizing the top 50 principal components (PCs) derived from the integrated dataset, we employed the Shared Nearest Neighbor approach to cluster the cells, resulting in 21 distinct clusters with a resolution set at 0.2. The R package ALDEx2 was employed to assess and compare the sizes of these clusters, as previously described (41).

### Preparation of BMDMs and LPMCs

BMDMs were generated following previously established protocols(42). For the isolation of colonic LPMC, the dissected mucosal tissue was incubated in calcium and magnesium-free Hank’s balanced salt solution (HBSS) (Life Technologies) containing 2.5% heat-inactivated fetal bovine serum (FBS) (Life Technologies) and 1 mM dithiothreitol (Sigma-Aldrich) to remove mucus. Epithelial cells were removed by incubation in HBSS containing 1 mM EDTA (Quality Biological) at 37°C for 30 min. After washing with HBSS, remaining tissues were collected and incubated, with agitation, in HBSS containing 400 U/ml type 3 collagenase and 0.01 mg/mL DNase I (Worthington Biochemical) for at 37°C 90 min. The digested cell fraction was pelleted, re-suspended in a 40% Percoll solution (GE Healthcare Life Sciences), layered on top of a 75% Percoll solution, and centrifuged at 700g for 20 min at room temperature. Viable LPMCs were recovered from the discontinuous gradient interface, as previously described(21, 22).

### Isolation of IL-1β-inducing pathobionts

To isolate commensal pathobionts with potent IL-1β-inducing capacity, colonic mucosal tissue was harvested from HMA mice colonized with CD or HC microbiota. The mucosal surface of the colonic tissue was scraped and homogenized in ice-cold, pre-reduced PBS within an anaerobic chamber. The mucosal homogenate was plated on various media (such as BHI with 5% FBS, GAM, Tryptic Soy, Schaedler with 5% horse blood, and Reinforced Clostridia with 5% horse blood) and cultured under anaerobic conditions. After 72 hrs, individual colonies were picked and further cultured in media (including BHI with 5% FBS, PYG, and Reinforced Clostridia media). The isolated bacterial species were identified by bacterial whole genome sequencing. To verify the IL-1β-inducing capacity of these isolated strains, BMDMs or colonic LPMCs were stimulated with isolated bacterial strains (at a MOI of 5) for 3 hrs. Gentamicin (100 μg/ml) was then added to prevent bacterial overgrowth, and the cells were incubated for an additional 16 hrs. The culture supernatants were collected, and cytokine levels were assessed using ELISA, as previously described(21, 22).

### Bacterial genomic analysis

Whole genome sequencing was conducted utilizing the Illumina MiSeq (500v2) system. High-quality paired-end reads, each 150 bp in length, were assembled into contigs through the use of SPAdes 3.13.0 (43), followed by annotation of the assembled contigs via Prokka(44). A phylogenetic tree was created for the new isolate HK alongside a reference *E. coli* strain, based on the presence (+) and absence (-) of all orthologues identified in the whole genome sequences using Roary after reannotation with Prokka (44), as previously described(45). In silico serotyping was carried out using BLASTN and BLASTP against the SeroTypeFinder O and H database(46), including K serotype markers such as KpsM, K1-specific KfoC, K2-specific Gtf1, alongside hypothetical proteins, and K3-specific KfoC, KfoG, K5-specific KfiA, and KfiB. Cut-off parameters were set to >80% identity and an e-value of <0.05.Orthologue groups were identified through Roary, while major virulence genes were identified via BLASTP against *E. coli* entries in the VFDB(47) and categorized based on VFDB descriptions.

### Bacterial adhesion and invasion assay

The T84 human colorectal carcinoma cell line was obtained from ATCC (Gaithersburg, MD) and grown in a mixture of Ham’s F-12 and DMEM (1:1), enriched with 10% FBS and an antibiotic solution (penicillin-streptomycin). The cells were plated at 2 × 105 density in 24-well plates for 21 days. Following this, the cells were exposed to bacteria at an MOI of 5-10, centrifuged at 1000g for 10 min, and incubated at 37°C for 3 hrs. For the adhesion assay, the cells underwent three PBS washes and were subsequently lysed using 0.1% Triton X-100 (Sigma-Aldrich) in deionized water. In the invasion assay, after a 3-hr incubation with bacteria, the cells were treated with gentamycin (100 µg/mL) for 1 hr to eliminate any extracellular bacteria. Afterward, the cells were washed three times with PBS and lysed using 0.1% Triton X-100 in deionized water. The lysates were diluted and plated on LB agar plates to quantify the CFUs related to the total cell-adhesive/invasive bacteria as previously described (22).

### IBIP-driven colitis model

SPF wild-type C57BL/6 mice were pre-treated with streptomycin and ampicillin (20 mg each) one day before bacterial inoculation. Following antibiotic administration, the mice were orally inoculated twice a week with either commensal *E. coli* MG1655 or IBIP *E. coli* HK7 at a dose of 1 × 10^9^ CFU per mouse. After 21 days of continuous bacterial colonization, the mice were given 1.5% DSS for 5 days, followed by regular water for an additional 2 days. To assess the involvement of the IL-1 pathway, one group of IBIP *E. coli* HK7-colonized mice received intraperitoneal injections of Anakinra (Swedish Orphan Biovitrum AB) 3 times per week (50 mg/kg per dose) from the start of bacterial colonization until day 28. Gut colonization by inoculated *E. coli* was confirmed by culturing fecal pellets on MacConkey agar plates containing 50 µg/ml streptomycin. Mice were euthanized on day 28 (seven days post-DSS), and colon tissues were harvested to assess inflammation. For histological analysis, colon tissues were fixed in 4% paraformaldehyde, processed, embedded, sectioned, and stained with H&E. Histological scores were assigned blindly by a trained pathologist based on the evaluation of the following variables(48, 49): severity of inflammation (0, none; 1, low density confined to mucosa; 2, moderate or higher density in mucosa and/or low to moderate density in mucosa and submucosa; 3, high density in submucosa and/or extension to muscular; 4, high density with frequent transmural extension), and extent of epithelial/crypt damage (0, none; 1, basal 1/3; 2, basal 2/3; 3, crypt loss; 4, crypt and surface epithelial destruction). Each variable was multiplied by a factor reflecting the percentage of the colon involved (1, 0%–25%; 2, 26%–50%; 3, 51%–75%; 4, 76%–100%). Each variable was then summed to obtain the overall score.

### Quantitative Real-Time PCR

RNA was extracted from colonic tissues using E.Z.N.A. Total RNA Kit I (Omega Bio-tek). RNA was reverse transcribed using a ReverTra Ace qPCR RT Master Mix (TOYOBO), and cDNA was then used for quantitative PCR analysis using PowerUp SYBR Green Master Mix (Thermo Fisher) on QuantStudio5 analyzer (Applied Biosystems). The following primer sets were used for amplification: *Actb*-F; 5’-AAGTGTGACGTTGACATCCG, *Actb*-R; 5’-GATCCACATCTGCTGGAAGG, *Il1b*-F; 5’-CAACCAACAAGTGATATTCTCCATG, *Il1b*-R; 5’-GATCCACACTCTCCAGCTGCA, *Il6*-F; 5’-GAGGATACCACTCCCAACAGACC, *Il6*-R; 5’-AAGTGCATCATCGTTGTTCATACA, *Tnf*-F; 5’-GCCTCCCTCTCATCAGTTCT, *Tnf*-R; CACTTGGTGGTTTGCTACGA.

### Microbiome analysis

Genomic DNA was isolated following an adapted protocol of the Qiagen DNeasy Blood and Tissue kit (Qiagen, Valencia, CA). The modifications comprised: [1] utilizing UltraClean fecal DNA bead tubes (Mo Bio Laboratories, Inc, West Carlsbad, CA) and a Mini-Beadbeater-16 (BioSpec Products, Inc, Bartlesville, OK) for sample homogenization (1.5 min); [2] increasing the initial buffer ATL volume from 180 to 400 µL; [3] doubling the volume of proteinase K from 20 to 40 µL; and [4] reducing the elution buffer AE volume from 200 to 75 µL at the protocol’s conclusion. The 16S rRNA sequencing was conducted by the University of Michigan Medical School Host Microbiome Initiative core facility using the MiSeq Illumina sequencing platform. Libraries for the 16S ribosomal RNA (rRNA) gene were generated with primers targeting the V4 region. The sequences were curated using the mothur software program (v.1.33)(50) and by adhering to the MiSeq SOP steps(51). Operational taxonomic units (OTUs) were identified with a 0.03 cutoff and classified using the Ribosomal Database Project (RDP) 16S rRNA gene training set (version 9) via a naïve Bayesian method, maintaining an 80% confidence threshold. The processed OTU sequence data were expressed as relative abundance ± standard error of the mean. Within-community diversity (α-diversity) was evaluated using the Shannon diversity index (H’) and OTU Richness. Between-community diversity (β-diversity) was assessed employing the Yue and Clayton (θYC) dissimilarity metric. Non-metric multidimensional scaling (NMDS) was applied to visualize the β-diversity results. An analysis of molecular variation (AMOVA) tested for significant structural differences with 10,000 permutations. To identify differentially abundant, biologically relevant functional pathways with notable effect sizes, we applied linear discriminant analysis effect size (LEfSe) as, previously described(7, 52).

### Measurement of fecal Lcn2 level by ELISA

Collected fecal samples were diluted 100-1000 fold with PBS. Lcn2 levels were determined in fecal supernatants using a Duoset murine Lcn2 ELISA kit (R&D Systems).

### Measurement of fecal IBIP-reactive IgA levels in humans

To measure IBIP-reactive IgA levels in fecal samples from HC individuals and CD patients, collected stool samples were diluted in PBS (100 mg/ml) and treated with a protease inhibitor (Sigma-Aldrich). The fecal suspension was then homogenized using a bead beater and centrifuged at 10,000 rpm for 5 min at 4°C. The supernatants were diluted 3-fold with PBS and passed through a 0.22 µm filter. For the detection of IBIP-reactive IgA, heat-inactivated IBIP strains HK7 or HK1 (2.5 × 10^7^ CFU in PBS) were coated onto a 96-well plate (Corning) and incubated overnight at 4°C. Following washes with PBST (PBS supplemented with 0.05% Tween 20), the plates were blocked with 5% BSA in PBS at room temperature for 1 hr. After blocking, plates were rinsed with PBST and incubated with the prepared stool samples (625 µg per well) for 1 hr at room temperature. After further washing with PBST, bacterial-specific IgA was detected using an HRP-conjugated goat anti-human IgA antibody (Southern Biotech). After additional washes with PBST, the plates were developed using TMB solution (Thermo Fisher), then the reaction was stopped with 2M sulfuric acid, and absorbance was measured at 450 nm using a microplate reader. Purified human IgA (Southern Biotech) served as the standard, and the cut-off value for high IgA was set as the mean plus three standard deviations(53).

### Pediatric Cohort

The Precision Crohn’s Disease Management Utilizing Predictive Protein Panels (ENvISION) trial (NCT04131504) was a multicenter, observational study of children and young adults with CD starting an anti-TNF biologic (infliximab or adalimumab). Baseline demographics, disease severity, and phenotype were collected while clinical, biochemical, endoscopic, and radiologic outcomes were monitored for up to one year or until the biologic was stopped or a patient had a CD-related surgery. The research-only stool was collected prior to starting anti-TNF with clinical remission determined by the weighted pediatric CD activity index (wPCDAI) and endoscopic severity assessed with the simple endoscopic score-CD (SES-CD). The study protocol was approved by the Institutional Review Board at all four participating centers, which included Cincinnati Children’s Hospital Medical Center, Connecticut Children’s, Nationwide Children’s, and the Medical College of Wisconsin.

### Statistical Analyses

Statistical analyses were performed using GraphPad Prism software version 10 (GraphPad Software Inc.). Statistical tests used for the analysis of data are identified in the legend of each figure. Differences of *P* < 0.05 were considered significant.

