## Supplementary material for "Immune phenotype-guided identification of disease-associated pathobionts in Crohn’s disease": Fig S1-S2, Table S1, S2

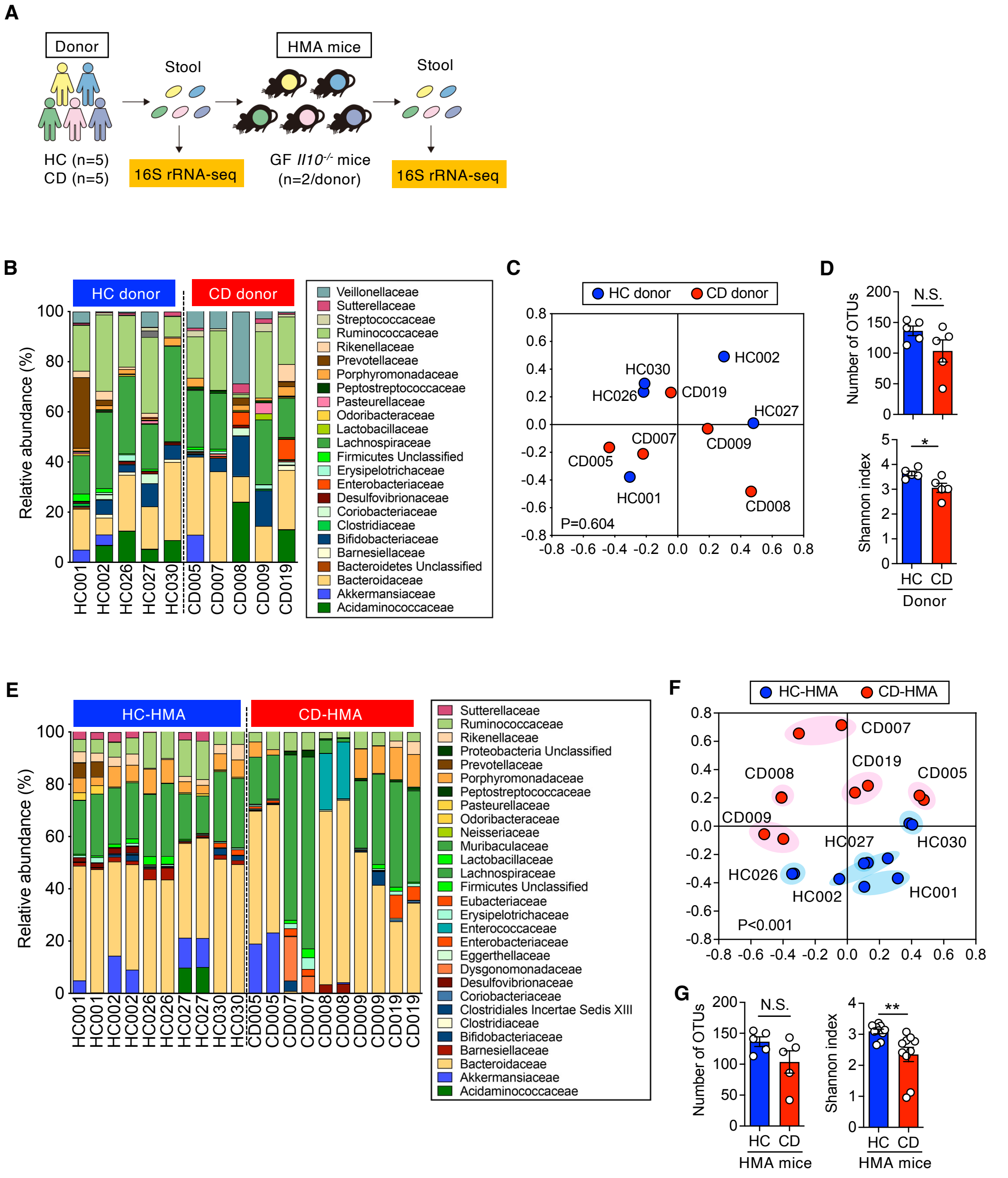


**Figure S1. Microbiome analysis of Crohn's disease donors and HMA mice.**

(A) The stool samples were obtained from 5 healthy control (HC) subjects and 5 patients with Crohn's disease (CD). Stool microbiotas were reconstituted in germ-free (GF) *Il10*$^{-/-}$ mice for 3 weeks to generate human microbiota-associated (HMA) mice (n=2 per donor). The gut microbiomes of both donors and HMA mice were analyzed using 16S rRNA sequencing. (B-G) Microbiome analysis: Relative bacterial abundance in donors (B) and HMA mice (E). Microbial community structures of donors (C) and HMA mice (F) were analyzed using the Yue & Clayton dissimilarity distance metric (θYC) and visualized in a non-metric multidimensional scaling (NMDS) plot. The number of operational taxonomic units (OTUs, richness) (left) and Shannon index (α-diversity) (right) in donors (D) and HMA mice (G) were assessed. Data are presented as mean ± SD. Dots represent individual patients or mice. N.S.: Not significant, *; $P < 0.05$**; $P < 0.01$ by Mann-Whitney *U* test.

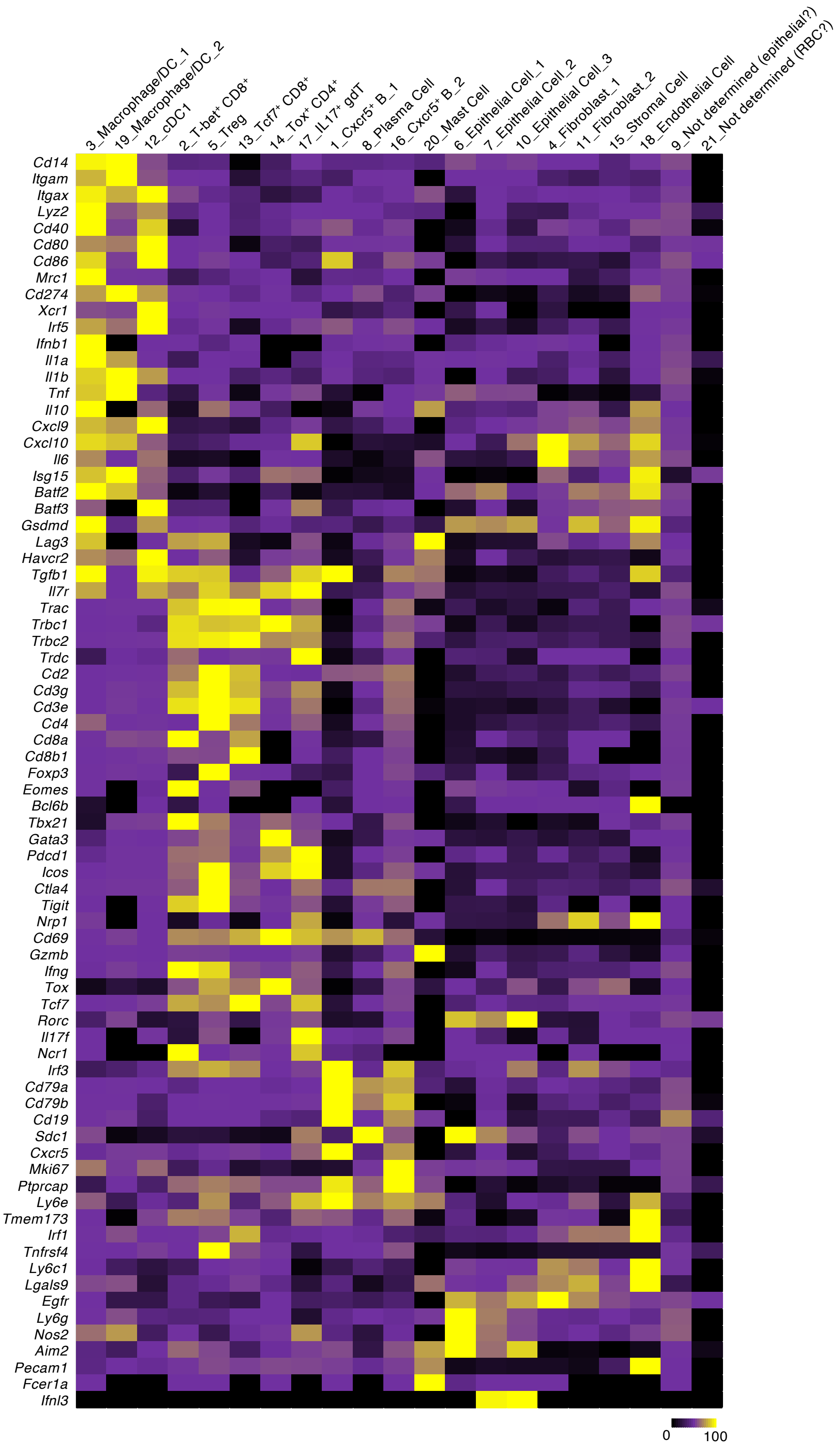


**Figure S2. Expression of genes used for cluster definition in scRNA-Seq analysis**
Relative expression levels of the indicated genes across 21 cell clusters.

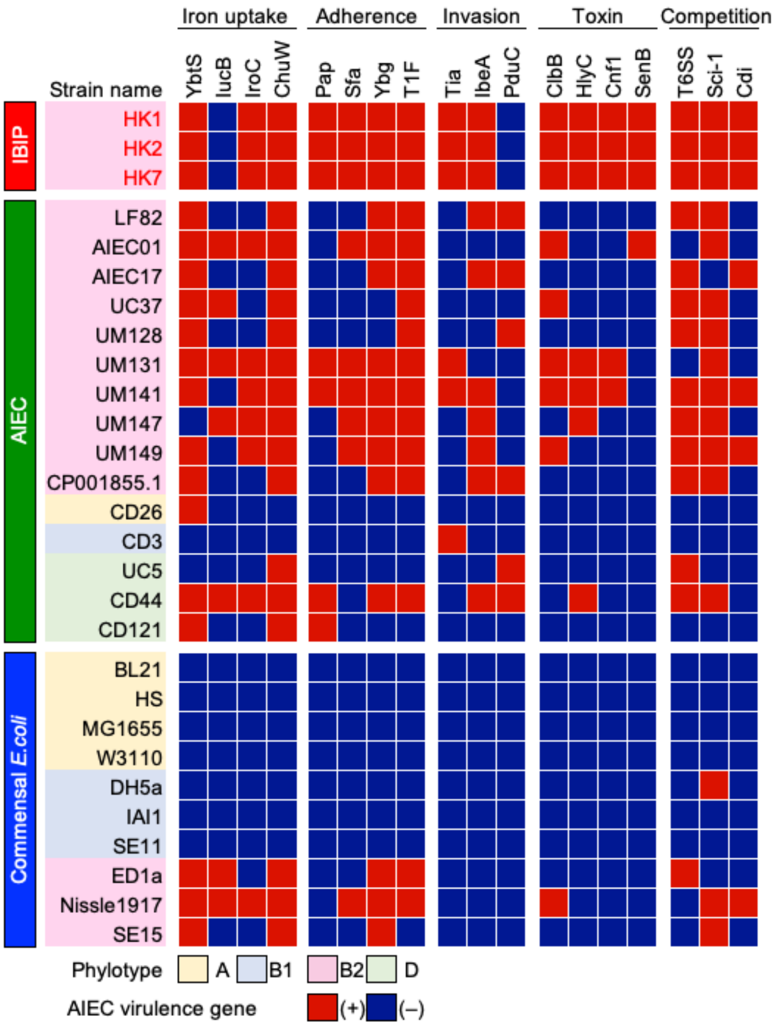


**Figure S3. Genetic characteristics of isolated IL-1β-inducing pathobionts.**
The heatmap illustrates the presence (red) or absence (blue) of orthologs associated with adherent-invasive *Escherichia coli* (AIEC) and extraintestinal pathogenic *Escherichia coli* (ExPEC) virulence genes.

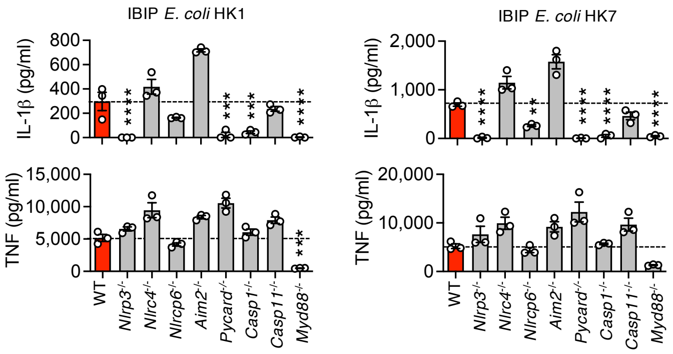


**Figure S4. Inflammasome-mediated secretion of IL-1β from macrophages upon IBIP stimulation.**

Bone marrow-derived macrophages (BMDMs) from wild-type, *Nlrp3*$^{-/-}$, *Nlrc4*$^{-/-}$, *Nlrp6*$^{-/-}$, *Aim2*$^{-/-}$, *Pycard*$^{-/-}$, *Casp1*$^{-/-}$, *Casp11*$^{-/-}$, and *Myd88*$^{-/-}$ mice were stimulated with IBIP *E. coli* strains HK1 or HK7. Secreted IL-1β and TNF levels were measured in the culture supernatant. Data are presented as mean ± SD. Dots represent biological replicates (n=3). *; $P < 0.05$, **; $P < 0.01$, ***; $P < 0.001$, ****; $P < 0.0001$, and by 1-Way ANOVA followed by Bonferroni post hoc test.

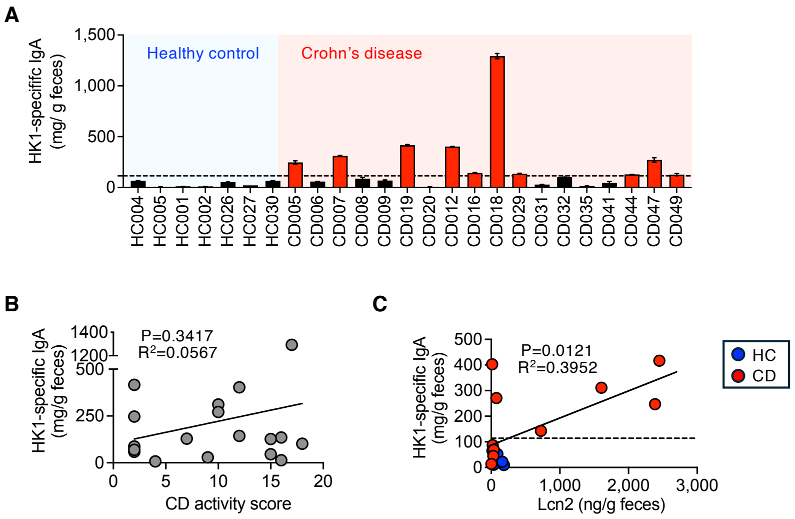


**Figure S5. IBIP-reactive IgA as a potential biomarker for predicting the colitogenic potential of the microbiota in CD patients**

(A) Levels of IBIP HK1-specific IgA in fecal samples from 7 HC and 18 CD patients. Data are presented as mean ± SD (*n* = 2 technical replicates). The dotted line represents the cut-off value (mean + 3 SD of HC samples). (B) Correlation between IBIP HK1-reactive IgA levels and CD activity scores (Harvey-Bradshaw Index, HBI). Statistical significance was determined using Pearson correlation analysis (two-sided). (C) Correlation between IBIP HK7-reactive IgA levels and the colitogenic capacity of microbiotas. The colitogenic potential of microbiotas was assessed by colonizing GF *Il10*$^{-/-}$ mice (HMA mice). After 3 weeks of bacterial reconstitution, fecal Lcn2 levels were measured as a marker of colitis. Statistical significance was determined using Pearson correlation analysis (two-sided).

**Table S1. Characteristics of adult Crohn's disease patients included in this study**

| Patient ID | Gender | Age | Race | Disease status | Disease location | CD activity score (HBI) |
|---|---|---|---|---|---|---|
| CD005 | M | 28 | Caucasian | Mildly active | Ileum | 2 |
| CD006 | M | 37 | Caucasian | Inactive | Ileum | 2 |
| CD007 | F | 25 | Caucasian | Active | Ileum | 10 |
| CD008 | M | 23 | Caucasian | Inactive | Ileo-colonic | 2 |
| CD009 | M | 27 | Caucasian | Active | Colon | 2 |
| CD019 | M | 28 | Caucasian | Active | Ileum | 2 |
| CD020 | M | 70 | Caucasian | Active | Ileo-colonic | 4 |
| CD012 | F | 45 | Caucasian | Active | Ileum | 12 |
| CD016 | M | 28 | Middle Eastern | Active | Colon | 12 |
| CD018 | F | 45 | Caucasian | Active | Ileum | 17 |
| CD029 | M | 29 | Caucasian | Active | Ileum | 16 |
| CD031 | F | 56 | Caucasian | Active | Colon | 9 |
| CD032 | F | 35 | Caucasian | Mildly active | Ileo-colonic | 18 |
| CD035 | M | 34 | Caucasian | Active | Ileum | 16 |
| CD041 | M | 27 | Caucasian | Active | Ileo-colonic | 15 |
| CD044 | M | 21 | Caucasian | Active | Ileum | 7 |
| CD047 | F | 37 | Caucasian | Active | Ileo-colonic | 10 |
| CD049 | M | 22 | Caucasian | Active | Ileo-colonic | 15 |

M: Male, F: Female, HBI: Harvey–Bradshaw Index.

**Table S2. Characteristics of treatment-naïve pediatric Crohn's disease patients included in this study**

| Pt ID | Gender | Therapy-treated age (year) | Race | Disease status | Disease location | Type of anti-TNF treatment | Clinical remission_3M (wPCDAI <12.5) | Endoscopic healing_12M (SES-CD < 4) |
|---|---|---|---|---|---|---|---|---|
| 1 | M | 10.6 | African American | Inflammation | Colon | Infliximab | No | 6 |
| 2 | M | 18.3 | White | Inflammation | Ileo-colonic | Adalimumab | Yes | N/A* |
| 3 | M | 9 | White | Stricture | Colon | Infliximab | No | 0 |
| 4 | M | 15.5 | White | Inflammation | Ileum | Infliximab | Yes | 0 |
| 5 | F | 13.4 | White | Inflammation | Ileo-colonic | Infliximab | Yes | 0 |
| 6 | F | 18.5 | African American | Stricture | Ileum | Adalimumab | No | 4 |
| 7 | F | 13 | White | Inflammation | Ileum | Infliximab | No | 2 |
| 8 | M | 15.9 | White | Inflammation | Ileo-colonic | Infliximab | Yes | 1 |
| 9 | F | 7.3 | White | Inflammation | Colon | Infliximab | Yes | 1 |
| 10 | M | 8.2 | White | Inflammation | Ileo-colonic | Infliximab | No | 13 |

* The patient did not undergo endoscopy 12 months after anti-TNF treatment

M: Male, F: Female, wPCDAI: weighted pediatric Crohn's disease activity index, SES-CD: Simple endoscopic score for Crohn's disease
